# Limitations of the ABEGO representation: ambiguity between αα-corner and αα-hairpin

**DOI:** 10.1101/2021.04.13.439694

**Authors:** Koya Sakuma

**Affiliations:** SOKENDAI, The Graduate University for Advanced Studies, 38 Nishigonaka, Myodaiji, Okazaki, 444-8585, Japan; Institute for Molecular Science, 38 Nishigonaka, Myodaiji, Okazaki, 444-8585, Japan

## Abstract

ABEGO is a coarse-grained representation for polypeptide backbone dihedral angles. The Ramachandran map is divided into four segments denoted as A, B, E, and G to represent the local conformation of polypeptide chains in the character strings. Although the ABEGO representation is widely used in structural informatics and protein design, it cannot capture minor differences in backbone dihedral angles, which potentially leads to ambiguity between two structurally distinct fragments. Here, we show a nontrivial example of two local motifs that could not be distinguished by their ABEGO representations. We found that two well-known local motifs αα-hairpins and αα-corners are both represented as α-GBB-α and thus indistinguishable in the ABEGO representation, although they show distinct arrangements of the flanking α-helices. We also found that α-GBB-α motifs caused a loss of efficiency in the ABEGO-based fragment-assembly simulations for protein backbone design. Nevertheless, we designed amino-acid sequences that were predicted to fold into the target topologies that contained these α-GBB-α motifs. Our finding that certain local motifs bottleneck the ABEGO-based fragment-assembly simulations for construction of backbone structures suggests that finer representations of backbone torsion angles are required for efficiently generating diverse topologies containing such indistinguishable local motifs.

## Introduction

Proteins are polymers, and using idealized bond lengths and bond angles, the conformation of a polypeptide chain can be represented as a series of backbone dihedral angle triplets (φ, ψ, and ω) [1]. Provided that all peptide bonds have *trans* conformations with ω of approximately 180°, the two-dimensional plot of φ and ψ called the Ramachandran map can have sufficient information to specify the residue-wise conformations of a polypeptide chain. To construct coarse-grained representations of backbone conformations, the Ramachandran map can be divided into subsections to cluster similar backbone conformations into the same class. A widespread approach is to define a four-state representation dividing the map into four segments and assigning the single letters A, B, E, and G to the regions (Figure 1). Broadly, the A region corresponds to α-helices and the B region to β-strands. For regions with positive φ, the G region corresponds to the left-handed α-helix and the E region represents the remaining map. With an additional state O corresponding to the *cis*-conformation of the peptide bond, this five-state discrete representation can cover the conformational space of polypeptide chains in a coarse-grained manner. This five-state coarse-grained representation of the polypeptide chain conformation is termed the ABEGO representation, which was the main focus of the current study.

**Figure 1:**
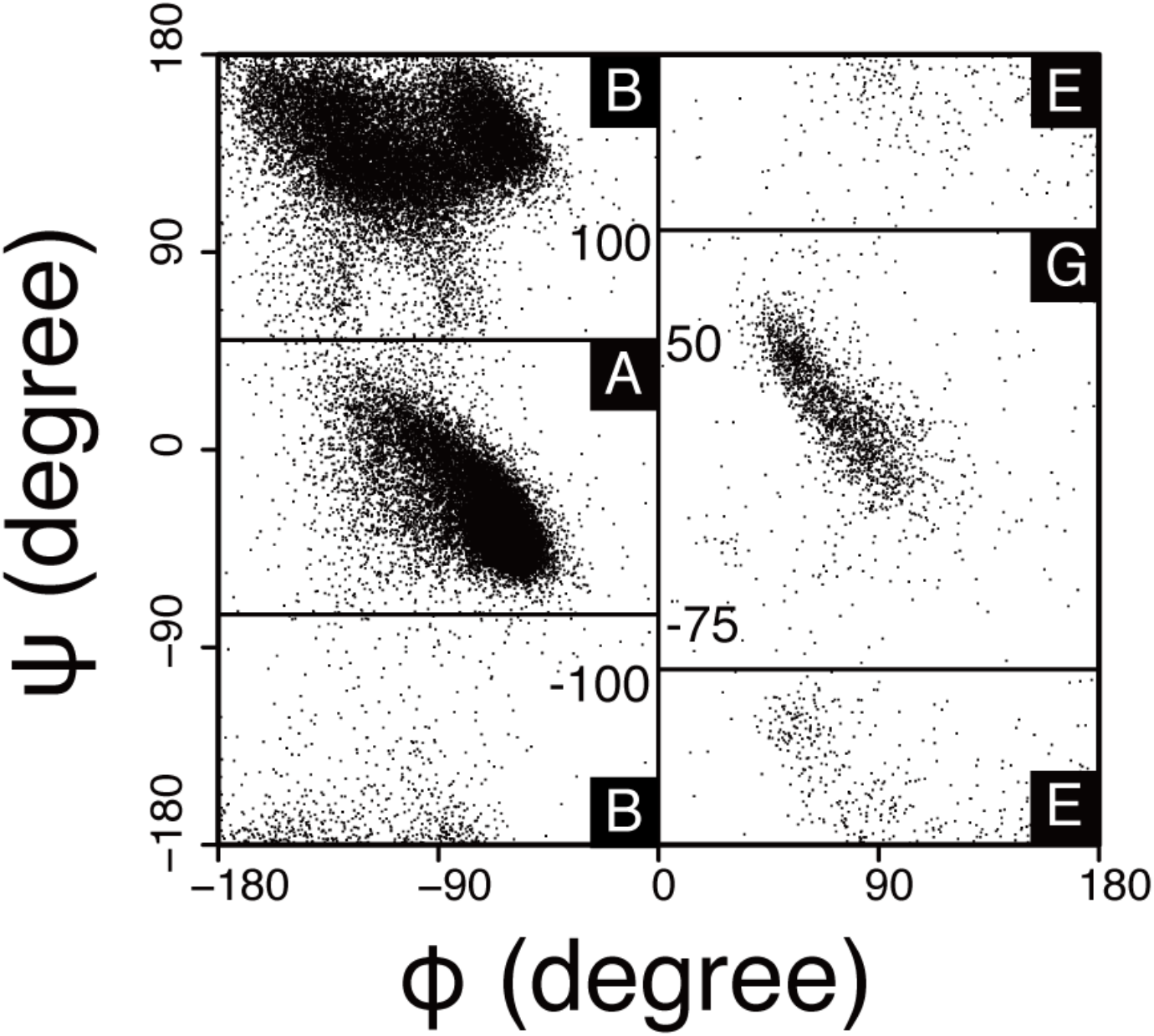
Definition of ABEGO. The horizontal axis represents φ, and the vertical axis represents the ψ angle of the polypeptide backbone structure. O is not defined in this diagram because it represents *cis*-peptides.

Because the ABEGO representation is useful for determining the conformation of short polypeptide fragments, it is routinely used in the classification of loop structures [2]. Another important application of the ABEGO representation is the *de novo* design of protein backbone structures. Designers specify the target topology using ABEGO sequences, select structure fragments that satisfy the desired ABEGO sequences, and perform fragment-assembly simulations to build the atomistic backbone structures with the desired topology [3]. Hereafter, we refer to these fragment-assembly simulations guided by ABEGO specification as ABEGO-based backbone-building simulations. This approach is widely accepted in *de novo* protein design and has been used to construct a variety of topologies ranging from small α-helical bundles to TIM barrels [4,5]. However, notably, ABEGO representations sometimes fail to distinguish two different conformations because it is a highly coarse-grained representation of backbone dihedral angles. In this study, we show non-trivial example of two famous local motifs that are indistinguishable by their ABEGO representation and point out that the ambiguity between these two motifs can lead to loss of efficiency in the ABEGO-based backbone building simulations. Clearifying the limiations of the ABEGO representation will motivate further development of more sophisticated representation for backbone confromation and backbone-building methods.

## Result

First, we investigated a nontrivial example in which ABEGO representation could not distinguish two structurally different local motifs. Using structural informatics analysis, we identified two distinct types of helix–loop–helix fragments that were indistinguishable based on their ABEGO sequences. Conformations of both motifs were represetanted α-GBB-α in their ABEGO representation, but they result in distinct overall structures and sequence preferences (Figure 2 and Figure S1). The first α-GBB-α motif is traditionally classified as an αα-corner that results in an almost orthogonal crossing angle between two flanking α-helices [6], and the second is called an αα-hairpin, which results in a steep hairpin turn for tightly packing adjacent α-helices into an antiparallel configuration [7]. Interestingly, their loop regions share nearly identical conformations; the residues on the C-terminal end of the N-terminal α-helix showed the most divergent conformations and were responsible for distinct overall arrangements of the two flanking α-helices (Figure S2). These two distinct conformations are identical in their ABEGO representation and are therefore indistinguishable in the coarse-grained representation.

**Figure 2:**
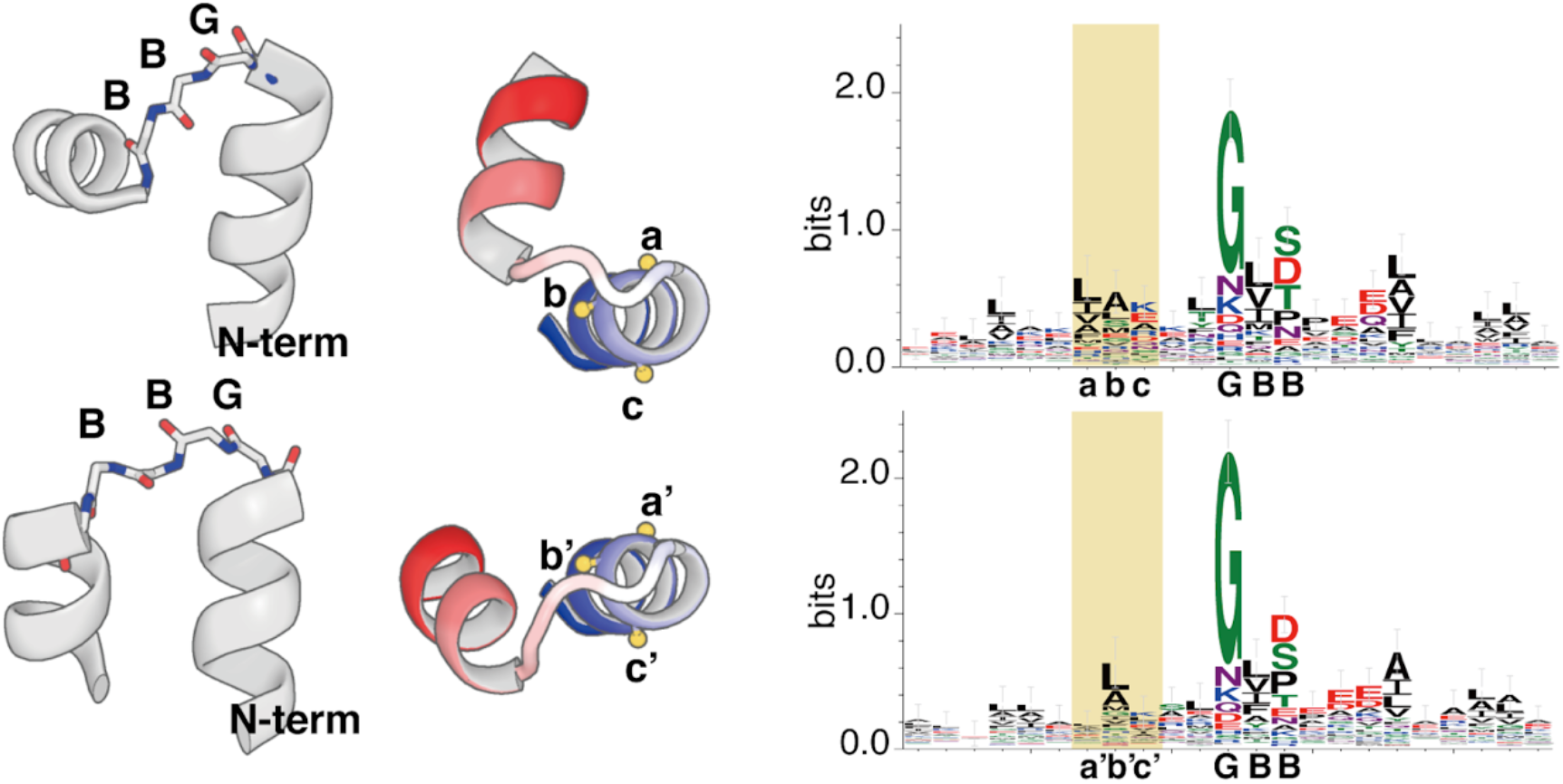
Comparison of αα-corner and αα-hairpins. These have similar backbone torsions but provide distinct contact patterns between two flanking α-helices. (Left) The overall structure of αα-corner and αα-hairpins. The loop regions are shown as sticks and colored in CPK scheme. The α-helices are shown in the cartoon representation. (Center) αα-corner and αα-hairpins offer different environments for nearby residues. Each fragment is colored in a blue–white–red gradient from the N-to C-terminus. The orange sphere represents the Cβ atoms on the N-terminal α-helical segments. The a in Cβ corresponds to a’, b to b’, and c to c’. Position a is more buried than position a’, and similarly, b is more exposed than b’. (Right) Sequence logo for αα-corner and αα-hairpins. The region shaded in orange corresponds to the residues whose Cβ atoms are colored in orange in the center panel. The alphabets beneath the logos indicate residue positions for the region on the N-terminus of loops, and the ABEGO backbone torsion angle representation for the loop regions. Because conformations remarkably differ between α-corner and αα-hairpins, the variance in the amino acid composition is most recognizable in the orange-shaded regions, which correspond to the flanking sequences, rather than in the loop region.

Next, we sought to identify whether the ambiguity between the αα-hairpin and αα-corner in the ABEGO representation causes severe loss of efficiency in ABEGO-based backbone-building simulations. Thus, we manually generated three types of four-helix up-down bundle structures: GBB, GB, and BAAB bundles (Figure 3). Based on these decoy structures, we specified the backbone dihedral angles in the ABEGO representations (Figure S3), selected the fragments satisfying the specification, and performed ABEGO-based backbone-building simulations [3,8]. Although the simulations for the GB and BAAB bundles successfully recovered the original four-helix up-down bundle topologies, we observed that the ABEGO-based backbone-building simulations for the GBB bundle failed to efficiently generate the target topology (Figure 3). In the simulations for the GBB bundle, most trajectories were trapped in misfolded structures that contained GBB corner fragments (Figure S4), which is undesirable for building the up-down bundles. Our results showed that the ABEGO specification was too ambiguous to restrict the fragment conformation and failed to separate the two types of α-GBB-α motifs. The inability of the ABEGO representation to separate these distinct structures caused the resultant fragment library to contain both αα-corners and αα-hairpins, and the mixture of these two α-GBB-α fragments lowered the sampling efficiency for the target four-helix up-down topology.

**Figure 3:**
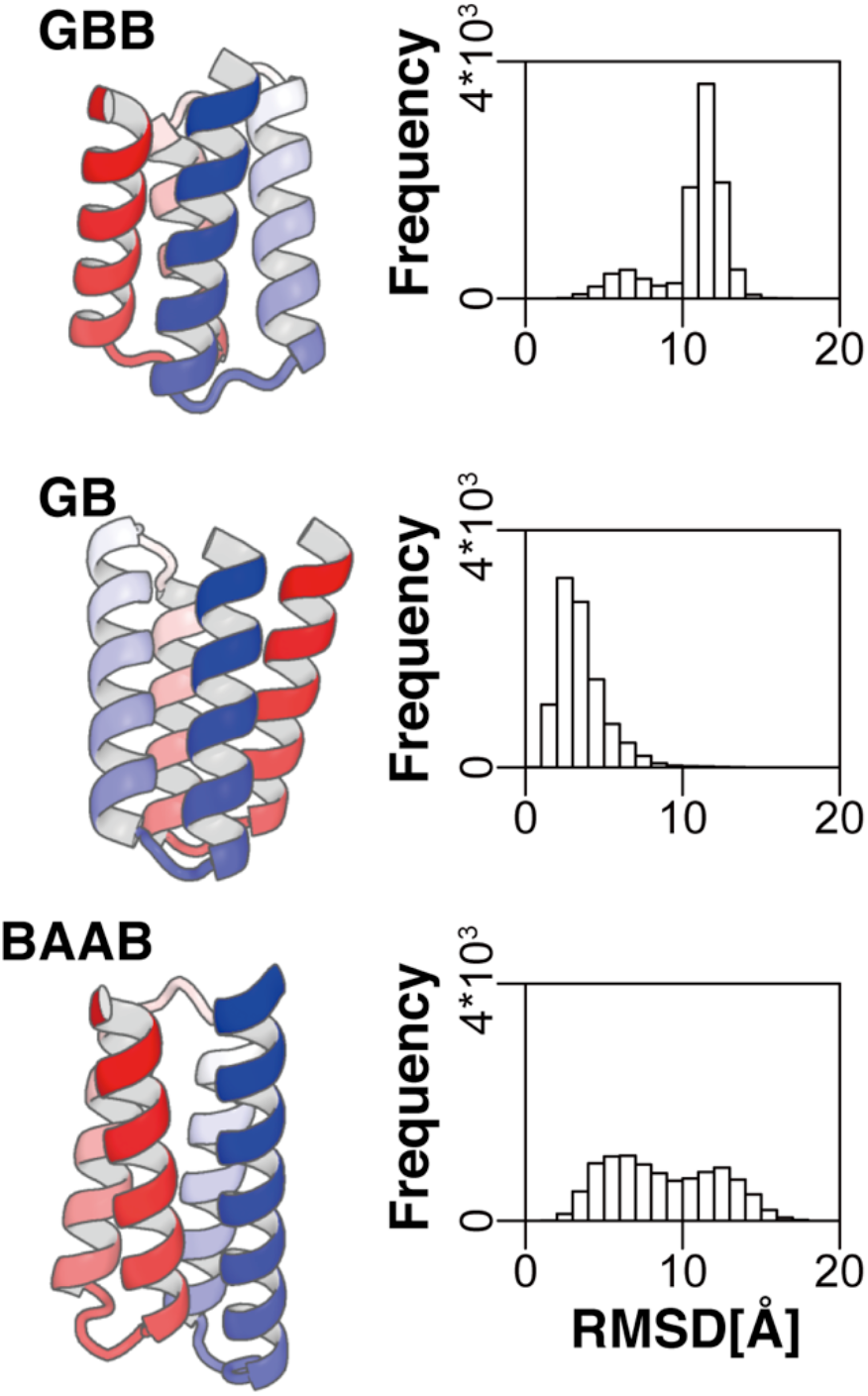
The foldability of four-helix up-down bundles in ABEGO-based backbone-building simulations. (Left) The structure of three types of four-helix up-down bundles. The ABEGO of hairpins are indicated on the left-top of each structure. (Right) The distribution of Cα RMSDs from 105 trajectories of ABEGO-based backbone-building simulations. The GBB bundle has a large peak around 10 Å, which indicates the ABEGO specification cannot force the polypeptide chain to fold into the target structure. The GB and BAAB bundles show reasonably large populations on the left (Cα RMSD < 5 Å), indicating that their ABEGO specification can allow the chain to fold into the target topology.

Considering the structures containing α-GBB-α fragments are difficult to compose in ABEGO-based backbone-building simulations, we sought to identify whether they can be designed when their amino acid sequences are completely specified. We performed an amino acid sequence design of two distinct structures composed of α-GBB-α motifs alone using Rosetta [8]. The first was the four-helix up-down bundle that we targeted to build as previously described, and the second was a small four-helix orthogonal bundle composed of two αα-hairpins and an αα-corner (Figure 4A and 4B). Similar to the ABEGO-based backbone-building simulations for the GBB up-down bundle, those for the GBB orthogonal bundle were also trapped in a misfolded state and showed low efficiency for achieving the target conformation (Figure S5), which supports the interpretation that α-GBB-α units lead to ambiguity during conformational sampling in ABEGO-based backbone-building simulations. However, by carefully designing amino acid sequences onto these structures using Rosetta, amino acid sequences that are predicted to fold into the respective target topologies can be obtained (Figure 4C and 4D). In contrast to the misfolding observed in the ABEGO-based backbone-building simulations, sequence-dependent fragment-assembly simulations successfully predicted both target topologies as having the lowest energy structures [9]. The results showed that plausible amino acid sequences can be designed once the backbone structures are built by some means even if they contain two indistinguishable α-GBB-α motifs. This result indicated that the conformational space that can be covered by the amino acid sequence design is broader than the conformational space in which ABEGO-based backbone-building simulations can firmly sample. Further, a novel backbone-building methodology may be required to improve the ability to generate more diverse and complicated backbone structures.

**Figure 4:**
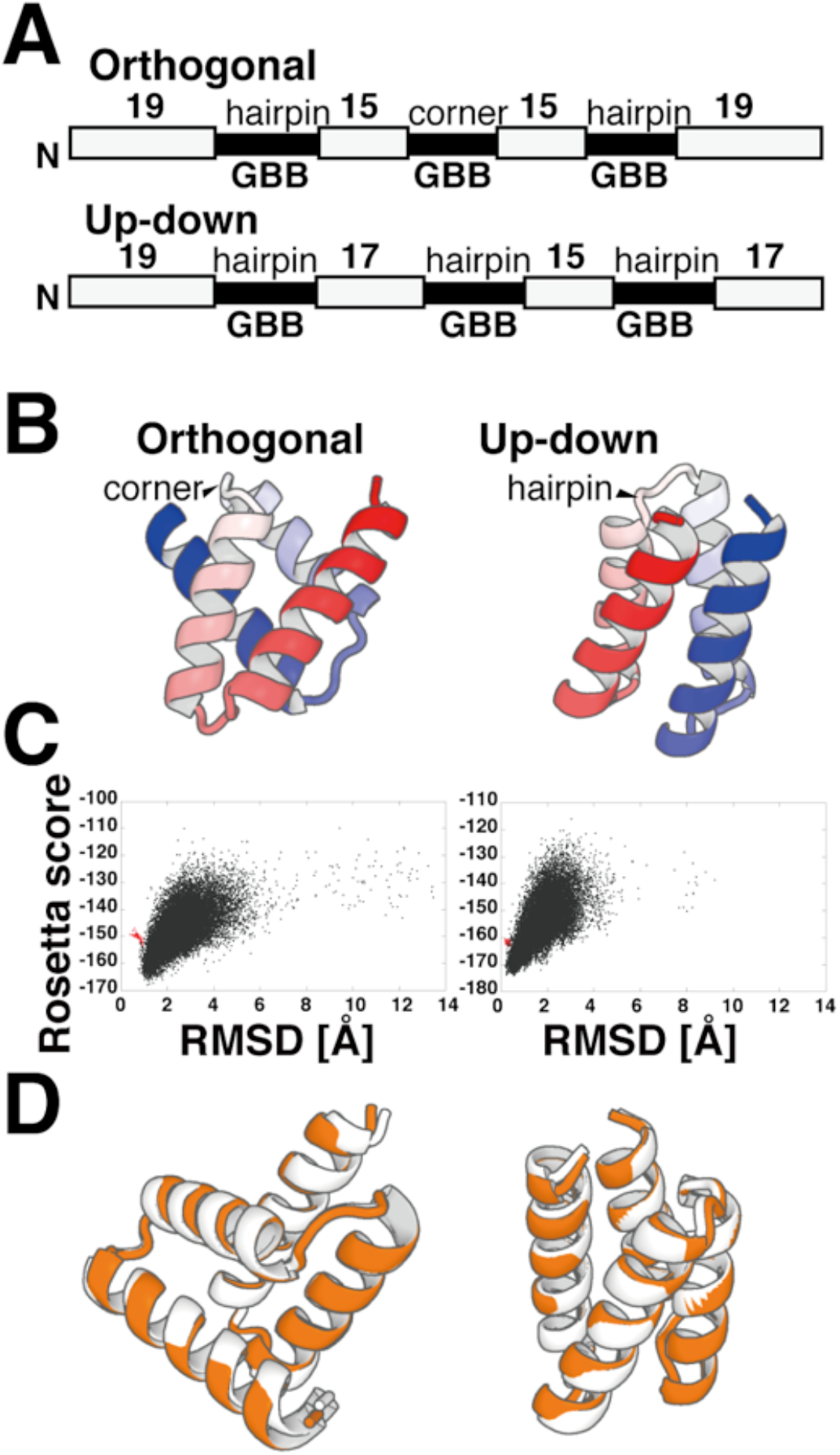
Design and sequence-dependent folding simulations of four-helix orthogonal and up-down bundles. (A) (B) Blueprints and structures of the GBB orthogonal bundles (left) and the up-down bundles (right). The gray bars represent the α-helices and black bars represent loop regions. As the loops are shown as GBB in the ABEGO representation, their intended structure types are indicated above the loop regions. (C) Energy-RMSD scatterplot from sequence-dependent folding simulations for orthogonal (left) and up-down bundles (right). Both designs have funneled energy landscapes and are predicted to fold into the target topology. (D) The superposition of the lowest energy structure (orange) onto the target structures. The lowest-energy structure from folding simulations of the orthogonal bundles showed Cα RMSD = 1.1 Å from the target conformation The lowest-energy structure for the up-down bundle showed Cα RMSD = 0.5 Å from the native.

## Conclusion

In this study, we showed that ABEGO is a coarse representation that can fail to distinguish different conformations, causing inefficiency in ABEGO-based backbone building for *de novo* protein design. The αα-corner and αα-hairpins are indistinguishable in the ABEGO representation because both are represented as α-GBB-α fragments. This ambiguity between these two distinct structures leads to difficulty in constructing simple four-helix bundle topologies composed of these α-GBB-α motifs. However, once a backbone structure is modeled by some means, amino acid sequences that specifically fold into the αα-corner and αα-hairpin can be designed. This enabled the selective design of two distinct types of four-helix bundle topologies.

Although we used the two indistinguishable α-GBB-α fragments as a nontrivial example in this study, such nondiscrimination may additionally occur for other motifs if the backbone torsion angles are represented in a coarse-grained manner. The apparent difficulty in designing structures that comprise indistinguishable local substructures lowers the efficiency of designing broad classes of protein structures and can be overcome by improving the methods used to construct backbone structures. Sequence design for such backbone structures does not appear to be very difficult, and we showed that two types of four-helix bundles composed of GBB fragments can be designed to fold into the target topologies. Therefore, novel methodologies for backbone building that can sample diverse structures unreachable by conventional structural modeling techniques will enable the design of a wide variety of protein structures. This will allow protein designers to further explore the protein structure universe and expand their design repertoires.

## Materials and methods

### Analysis of helix–loop–helix fragments

We composed a set of 29,397 non-redundant domain structures, which were a subset of the Evolutionary Classification Of protein Domains database (version develop238) culled by 40% sequence identity [10]. Structures with residues less than and equal to nine were discarded. Next, secondary structures were assigned using the DSSP [11]. The ABEGO representations of backbone torsion were assigned using in-house Python scripts according to the definition shown in Figure 1, and the fragments with GBB loops were extracted. We calculated the all-to-all Cα root mean square deviation (RMSD) within these GBB fragments and performed k-medoid clustering with k = 2. Next, we defined the helix–helix crossing angle (Figure S1) using the helix orientation vector defined by Krissinel et al. [12] and confirmed that the clustering can clearly separate αα-corners and αα-hairpins.

### Construction of target structures

We composed the GBB, GB, and BAAB up-down bundles as well as the GBB orthogonal bundle by manually grafting the helix–loop–helix fragments using PyMOL (The PyMOL Molecular Graphics System, version 2.0 Schrödinger, LLC.) and removed severe steric clashes using Foldit [13]. The constructed backbone structures were used as templates for the ABEGO specifications, and the reference structures for the ABEGO-based backbone-building simulations. These structures were also used as template backbones for amino acid sequence design by Rosetta.

### Backbone-building simulations

Sequence-independent fragment assembly simulations, termed ABEGO-based backbone-building simulations, were performed using Rosetta BluePrintBDR [8], as described by Lin et al. [3]. Blueprint files were generated based on the target backbone structure that was manually built in advance, and the files were used for fragment selection to specify the backbone torsion in the ABEGO representation. For each ABEGO specification, simulations were repeated for 10,000 trajectories, and the final snapshots from the trajectories were used for structural analysis. During the analysis, we calculated the Cα RMSD of each structure referenced by the target backbone structures.

### Amino acid sequence design and sequence-dependent folding simulations

We performed amino acid sequence designs using the Rosetta flxbb protocol [8] starting from the backbone structure that was built manually. To enhance the efficiency of sequence design, amino acid profiles were constructed for the loop region using similar loop structure fragments (Cα RMSD < 2 Å) and were used as constraints for the residues used, as described by Marcos et al. [14]. The specifications of the residues were refined based on the buriedness of the backbone atoms using in-house programs. We performed 10,000 design trials for each backbone model, selected the best sequences based on the fragment-quality score, and performed sequence-dependent fragment-assembly folding simulations [9] to identify the best design sequences. We defined the fragment-quality score as the average of the logarithm of the number of fragments that had a Cα RMSD value lower than 1.5 Å in the design model. A total of 20,000 trajectories for folding simulations were obtained for each design protein to check the foldability.

## Acknowledgments

K.S. would like to thank Dr. Shintaro Minami for providing a curated domain structure dataset and the Koga laboratory for offering computational resources. Most of the computational analysis was performed using the facilities at the Research Center for Computational Science, Okazaki, Japan. K.S. was supported by a Grant-in-Aid for JSPS Fellows (grant number 15J02427). Additionally, K.S. would like to thank the Institute for Molecular Science for the financial support received as a research assistant during the doctoral course.

## Author Contributions

K.S. designed the research, performed numerical experiments, analyzed data, and wrote manuscripts.

---

**Figure S1:**
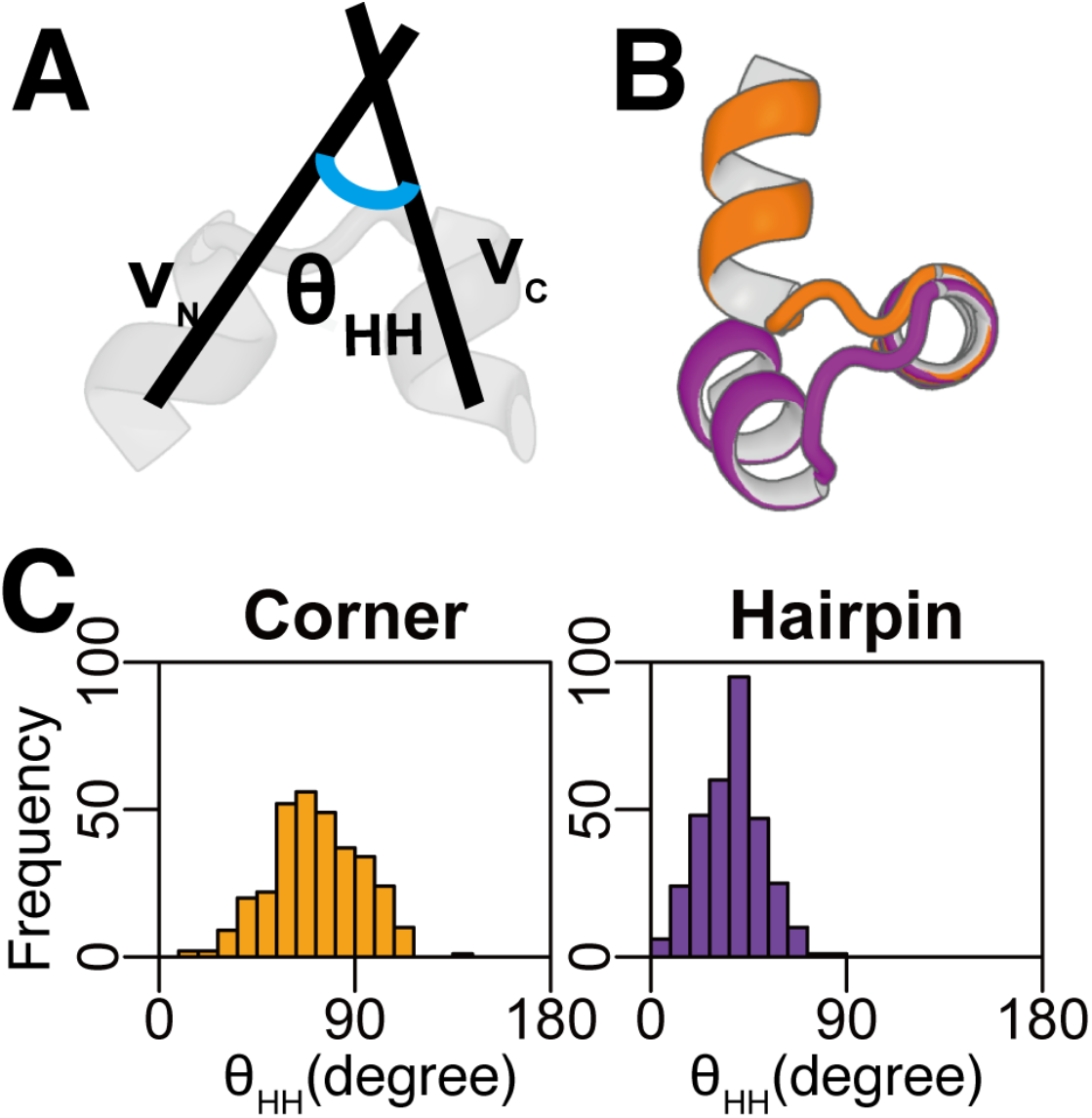
Comparison of αα-corners and αα-hairpins based on the helix-helix crossing angles. αα-corner and αα-hairpins are easily distinguished by helix-helix crossing angles. (A) Definition of helix-helix crossing angle. (B) αα-corner (orange) and αα-hairpins (purple) are superimposed to show their distinct conformations. Both structures are cluster representatives (medoids) derived by k-medoids clustering with k=2 when considering the all-to-all Cα RMSDs as the metric. (C) The distribution of helix-helix crossing angles after k-medoids clustering with k=2. The αα-corner and αα-hairpins have a distinct distribution of helix-helix crossing angles.

**Figure S2:**
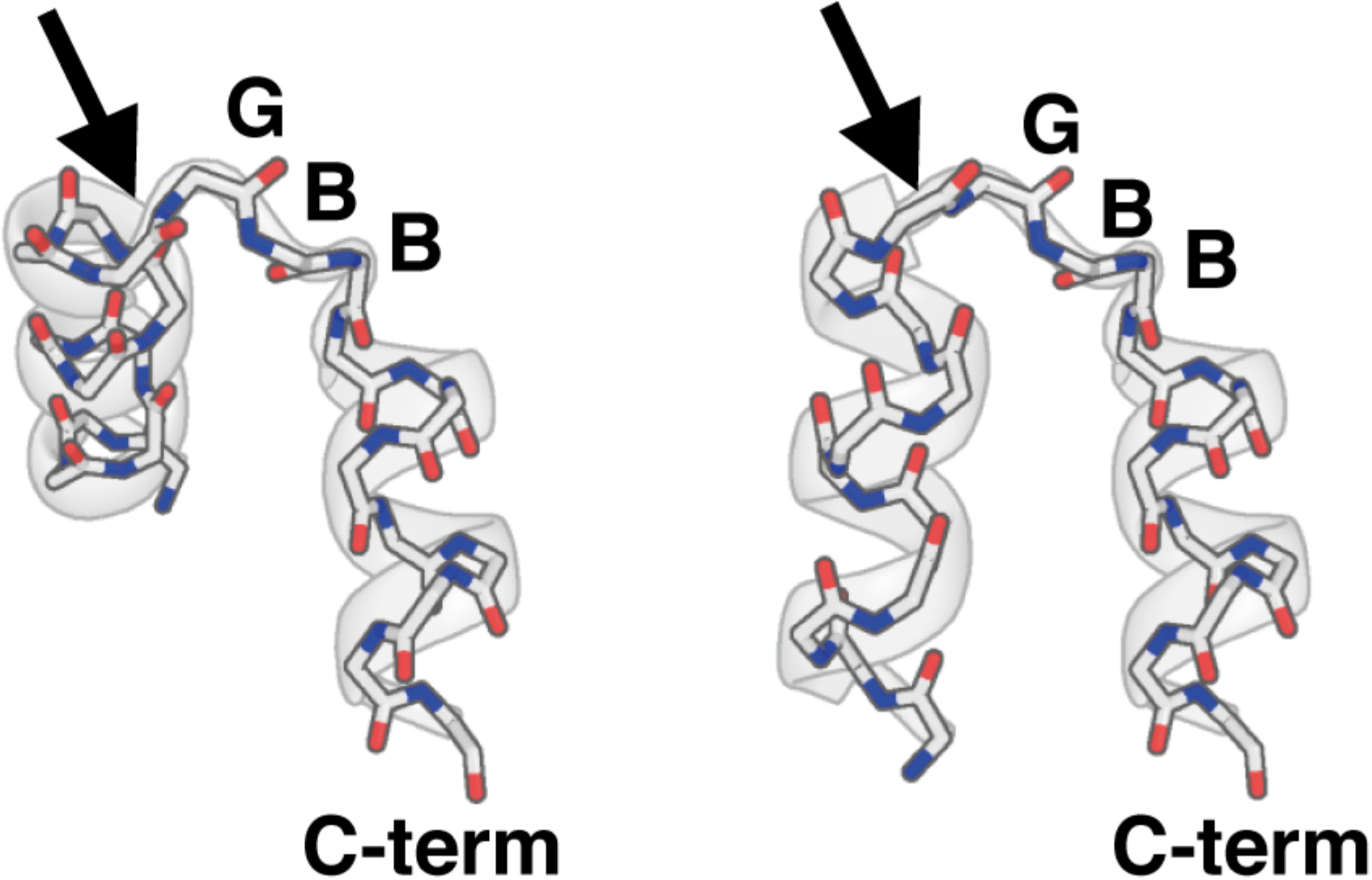
The residue responsible for the large deviation between αα-corner (left) and αα-hairpin (right). The backbone torsion angles of loop residues are indicated by ABEGO. The arrows point to the final residue of the N-terminal α-helix, which is responsible for the differentiation between αα-corners and αα-hairpins.

**Figure S3:**
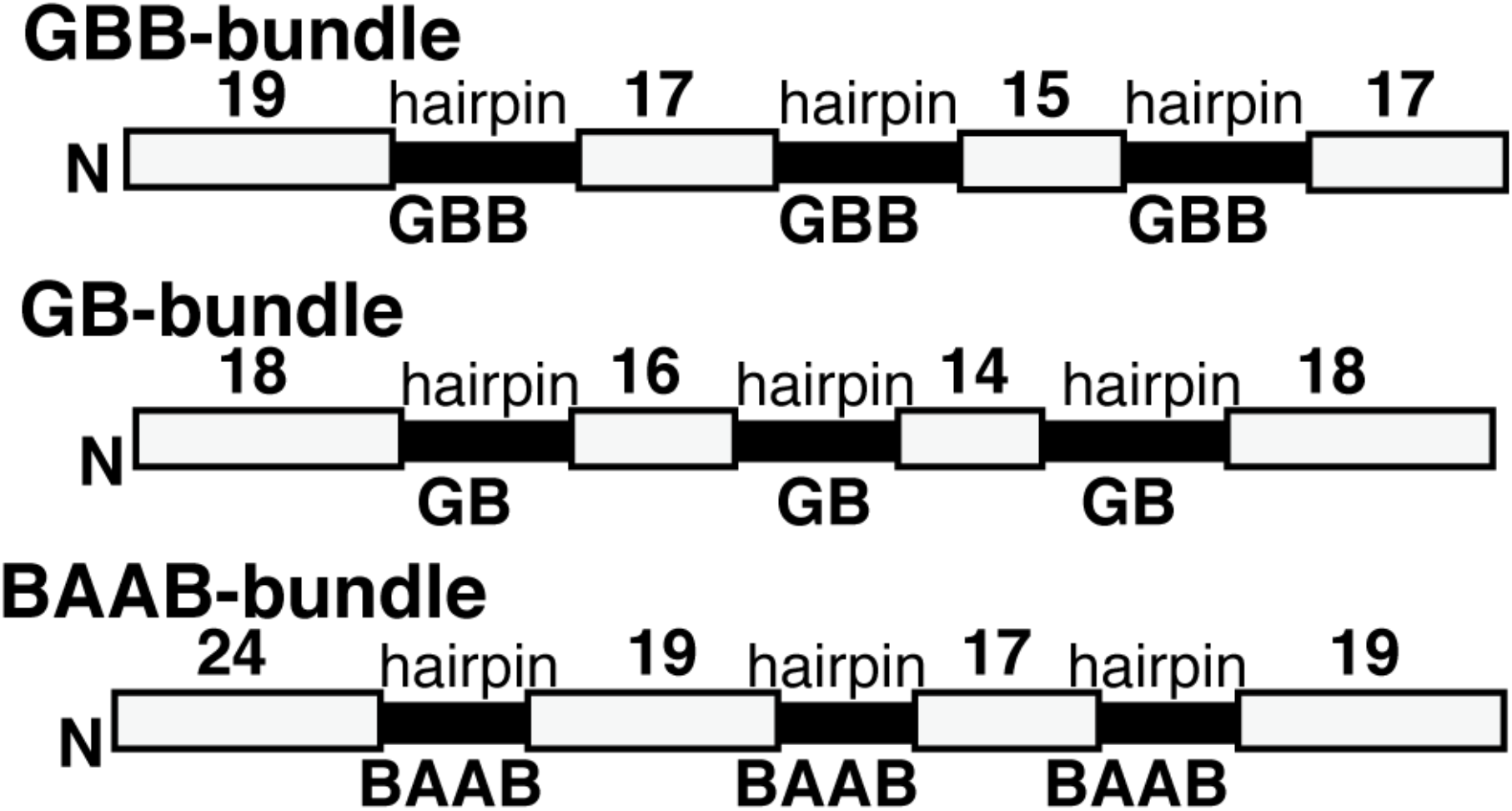
Blueprints for GBB, GB, and BAAB four-helix up-down bundles. The grey bars represent the α-helices and the black bars represent loop regions. The ABEGO of the loop is indicated beneath the loop region. The numbers indicate the residue number in the α-helix.

**Figure S4:**
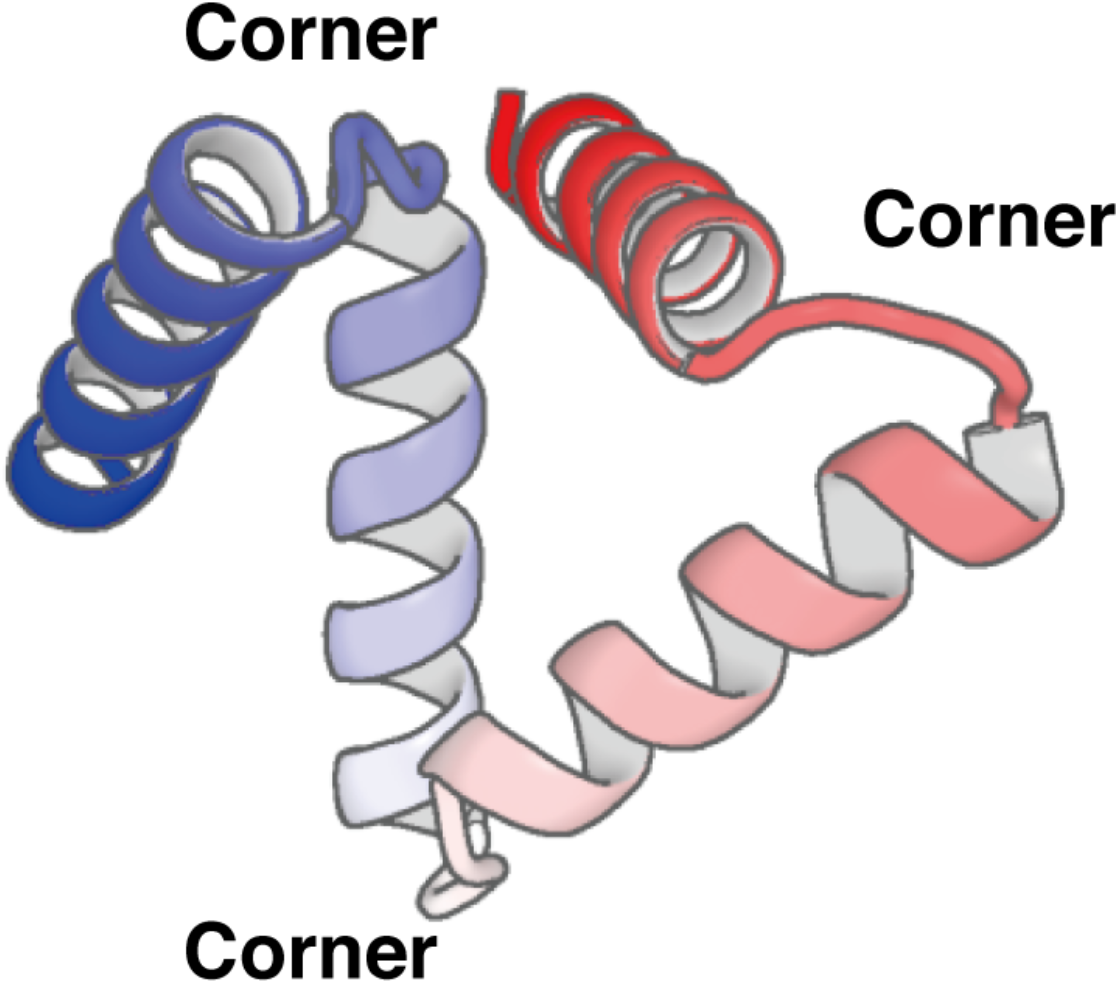
An example of a misfolded structure observed in ABEGO-based backbone-building simulations for the GBB up-down bundle. The loop regions take αα-corner conformations instead of αα-hairpins. This ambiguity between corners and hairpins makes it difficult to construct a simple up-down bundle structure by ABEGO-based backbone-building simulations.

**Figure S5:**
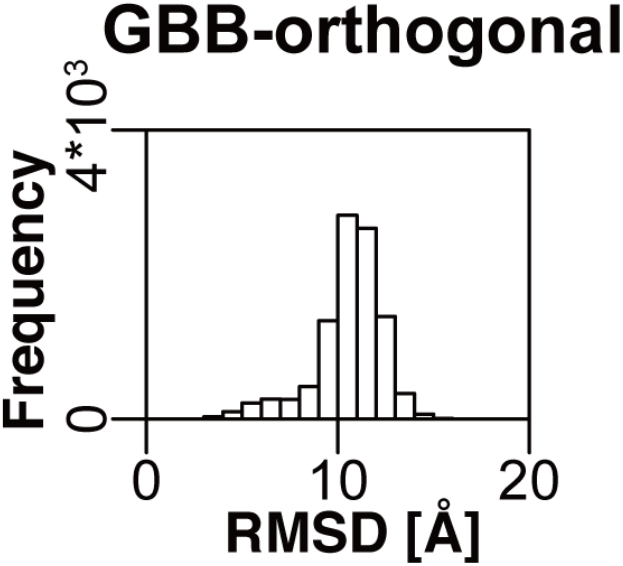
Distribution of RMSDs from the sequence-independent ABEGO-based backbone building simulations of four-helix orthogonal bundles. Similar to the four-helix up-down bundle composed of GBB-hairpins, the distribution of the GBB-orthogonal bundles also showed a large peak around 10 Å. This indicates the GBB orthogonal bundle is difficult to build using ABEGO-based backbone building simulations.

## Notes

### Competing Interest Statement

The authors have declared no competing interest.

## References

[1] G.N. Ramachandran, C. Ramakrishnan, V. Sasisekharan, Stereochemistry of polypeptide chain configurations, J. Mol. Biol. 7 95–99 (1963). DOI: 10.1016/S0022-2836(63)80023-6.

[2] R.T. Wintjens, M.J. Rooman, S.J. Wodak, Automatic classification and analysis of αα-turn motifs in proteins, J. Mol. Biol. 255 235–253 (1996). DOI: 10.1006/jmbi.1996.0020.

[3] Y.R. Lin, N. Koga, R. Tatsumi-Koga, G. Liu, A.F. Clouser, G.T. Montelione, D. Baker, Control over overall shape and size in de novo designed proteins, Proc. Natl. Acad. Sci. U. S. A. 112 E5478–E5485 (2015). DOI: 10.1073/pnas.1509508112.

[4] G.J. Rocklin, T.M. Chidyausiku, I. Goreshnik, A. Ford, S. Houliston, A. Lemak, L. Carter, R. Ravichandran, V.K. Mulligan, A. Chevalier, C.H. Arrowsmith, D. Baker, Global analysis of protein folding using massively parallel design, synthesis, and testing, Science 357 168–175 (2017). DOI: 10.1126/science.aan0693.

[5] P.S. Huang, K. Feldmeier, F. Parmeggiani, D.A.F. Velasco, B. Höcker, D. Baker, De novo design of a four-fold symmetric TIM-barrel protein with atomic-level accuracy, Nat. Chem. Biol. 12 29–34 (2016). DOI: 10.1038/nchembio.1966.

[6] A.V. Efimov, A novel super-secondary structure of proteins and the relation between the structure and the amino acid sequence, FEBS Lett. 166 33–38 (1984). DOI: 10.1016/0014-5793(84)80039-3.

[7] A.V. Efimov, Structure of α-α-hairpins with short connections, Protein Eng. Des. Sel. 4 245–250 (1991). DOI: 10.1093/protein/4.3.245.

[8] S.J. Fleishman, A. Leaver-Fay, J.E. Corn, E.M. Strauch, S.D. Khare, N. Koga, J. Ashworth, P. Murphy, F. Richter, G. Lemmon, J. Meiler, D. Baker, Rosettascripts: A scripting language interface to the Rosetta Macromolecular modeling suite, PLOS ONE 6 e20161 (2011). DOI: 10.1371/journal.pone.0020161.

[9] P. Bradley, K.M.S. Misura, D. Baker, Toward high-resolution de novo structure prediction for small proteins, Science 309 1868–1871 (2005). DOI: 10.1126/science.1113801.

[10] H. Cheng, R.D. Schaeffer, Y. Liao, L.N. Kinch, J. Pei, S. Shi, B.H. Kim, N.V. Grishin, ECOD: An evolutionary classification of protein domains, PLOS Comput. Biol. 10 e1003926 (2014). DOI: 10.1371/journal.pcbi.1003926.

[11] W. Kabsch, C. Sander, Dictionary of protein secondary structure: Pattern recognition of hydrogen◻bonded and geometrical features, Biopolymers 22 2577–2637 (1983). DOI: 10.1002/bip.360221211.

[12] E. Krissinel, K. Henrick, Secondary-structure matching (SSM), a new tool for fast protein structure alignment in three dimensions, Acta Crystallogr. D Biol. Crystallogr. 60 2256–2268 (2004). DOI: 10.1107/S0907444904026460.

[13] R. Kleffner, J. Flatten, A. Leaver-Fay, D. Baker, J.B. Siegel, F. Khatib, S. Cooper, Foldit Standalone: A video game-derived protein structure manipulation interface using Rosetta, Bioinformatics 33 2765–2767 (2017). DOI: 10.1093/bioinformatics/btx283.

[14] E. Marcos, B. Basanta, T.M. Chidyausiku, Y. Tang, G. Oberdorfer, G. Liu, G.V.T. Swapna, R. Guan, D.A. Silva, J. Dou, J.H. Pereira, R. Xiao, B. Sankaran, P.H. Zwart, G.T. Montelione, D. Baker, Principles for designing proteins with cavities formed by curved β sheets, Science 355 201–206 (2017). DOI: 10.1126/science.aah7389.

